# Normal gastric tissue *Helicobacter pylori* infection is associated with epigenetic age acceleration, increased mitotic tick rate, tissue cell composition, and Natural Killer cell methylation alterations

**DOI:** 10.1101/2023.06.28.546926

**Authors:** Irma M. Vlasac, Brock C. Christensen, Lucas A. Salas

**Affiliations:** Department of Epidemiology, Geisel School of Medicine at Dartmouth, Lebanon, NH 03756, USA; Department of Molecular and Systems Biology, Geisel School of Medicine at Dartmouth, Lebanon, NH 03756, USA

**Author notes:** Corresponding Author: Irma M. Vlasac.

**Keywords:** Gastric cancer, *H. pylori* infection, DNA methylation, epigenetics, epigenetic clocks, age acceleration, cell proportions, immune cells

## Abstract

**Background:** Gastric adenocarcinomas are a leading cause of global mortality, associated with chronic infection with *Helicobacter pylori*. The mechanisms by which infection with *H. pylori* contributes to carcinogenesis are not well understood. Recent studies from subjects with and without gastric cancer have identified significant DNA methylation alterations in normal gastric mucosa associated with *H. pylori* infection and gastric cancer risk. Here we further investigated DNA methylation alterations in normal gastric mucosa in gastric cancer cases (n = 42) and control subjects (n = 42) with *H. pylori* infection data. We assessed tissue cell type composition, DNA methylation alterations within cell populations, epigenetic aging, and repetitive element methylation.

**Results:** In normal gastric mucosa of both gastric cancer cases and control subjects, we observed increased epigenetic age acceleration associated with *H. pylori* infection. We also observed an increased mitotic tick rate associated with *H. pylori* infection in both gastric cancer cases and controls. Significant differences in immune cell populations associated with *H. pylori* infection in normal tissue from cancer cases and controls were identified using DNA methylation cell type deconvolution. We also found natural killer cell-specific methylation alterations in normal mucosa from gastric cancer patients with *H. pylori* infection.

**Conclusions:** Our findings from normal gastric mucosa provide insight into underlying cellular composition and epigenetic aspects of *H. pylori* associated gastric cancer etiology.

## Background

*Helicobacter pylori* has been established as a causative infectious agent in the development of gastric cancer and other gastric diseases. Chronic colonization of the gastric epithelia by *H. pylori* has an established pathogenic cascade(1); however, the molecular mechanisms by which *H. pylori* induces gastric cancer remain to be clearly elucidated(2). Gastric cancer is one of the leading causes of global mortality, accounting for 768,793 (7.7%) new cancer deaths in 2020 and 1,089,103 (5.6%) of newly diagnosed cancer cases(3). While infection with *H. pylori* is not sufficient for the development of gastric cancer, complete eradication of *H. pylori* lowers the risk of developing gastric cancer(4), indicating multiple additional factors are involved in gastric carcinogenesis. The interplay between *H. pylori* virulence factors, host immune response, and genetic and epigenetic alterations at the cellular level highlights the role of inflammation as a driver of gastric cancer(5). A recent study assessed the influence of *H. pylori* on inflammation and age-related methylation, marking age-related methylation differences between gastric mucosae from young and old subjects, as well as differences in DNA methylation specific to inflammation, further indicating the influence of *H. pylori* on methylation through infection and chronic inflammation(6).

DNA methylation is an epigenetic modification involved in normal development and cell differentiation, demonstrating unique methylation patterns by cell type. Alterations to DNA methylation in carcinogenesis contribute to genomic deregulation, genomic instability, and gene silencing(7). Genome-wide hypomethylation and hypermethylation of tumor suppressor gene loci are key characteristics of cancer cells, and aberrant DNAm contributes significantly to disease and disease susceptibility(8–10). Recent studies suggest that altered DNA methylation from *H. pylori* infection results from inflammation and inflammation-related processes rather than *H. pylori* infection alone(11). Altered DNA methylation may be cell type-specific, and through cell deconvolution of DNA methylation data, identifying specific cell types whose altered DNA methylation is associated with cancer is crucial to understanding drivers of carcinogenesis. Aberrant DNA methylation may also result in hypomethylation of sites typically hypermethylated and transcriptionally repressed, such as at repetitive elements, where hypomethylation of these sequences correlates with open chromatin and aberrant transcription, contributing to genomic instability(12). Additionally, since DNA methylation serves as a marker of tissue aging and cell division, it can be leveraged to calculate the difference between the DNA methylation age and chronological age (age acceleration)(13), and to estimate the cumulative number of stem cell divisions in a tissue (mitotic tick rate)(14). Cancer risk may correlate with the cumulative number of cell divisions in tissues, and assessing the mitotic age of normal gastric mucosa in the context of *H. pylori* infection may provide greater insight into how tissues may vary based on *H. pylori* infection(15). Assessing patterns of epigenome deregulation via DNA hypo- and hypermethylation provides a basis for understanding cancer. The epigenome serves as a marker of environmental exposures, allowing us to investigate how *H. pylori*-driven inflammation alters the gut cell microenvironment and epigenome and contributes to cancer pathogenesis.

Recent work by Woo et al. aimed to investigate how *H. pylori* affects DNA methylation in normal gastric mucosa from gastric cancer cases and controls by *H. pylori* infection status. Regions of the genome that are hypermethylated in association with *H. pylori* infection as well as DNA methylation signatures predictive of gastric cancer risk independent of *H. pylori* status were identified(16). We aimed to further study the impact of *H. pylori* infection on DNA methylation by assessing the impact on DNA methylation age, age acceleration, cell proportions, and methylation of genomic repetitive elements.

Here we demonstrate influence on epigenetic mitotic tick rate, changes to DNA methylation at repetitive elements, and differences in cell proportions based on *H. pylori* infection status in normal gastric mucosa from patients with and without gastric cancer.

## Methods

### Study population

Raw DNA methylation data IDAT files were downloaded from Gene Expression Omnibus (GEO) GSE99553, which included data from normal gastric mucosa from 84 subjects comprised of 42 gastric cancer cases and 42 controls. The 84 subjects were further stratified by *H. pylori* status of negative infection, a past infection, and positive infection, with case and control groups containing the same number of subjects per *H. pylori* status. Subject age data for GSE99553 was provided on request from the authors.

### Statistical Analysis

All data analysis was conducted in R version 4.1.2. Infinium HumanMethylation450K Beadchip Kit (HM450 array) IDAT files were preprocessed using minfi and ENmix.(17) Probes with out-of-band hybridization >0.05 were excluded from the analyses^10^, followed by normalization and background correction using ENmix. Probe type correction for HM450K was completed using beta-mixture quantile normalization (BMIQ)(18). Cross-reactive and polymorphic probes, SNP probes, and sex probes were masked in data analyses using Zhou annotation(19)

DNA methylation clocks were calculated using ENmix methyAge, including Horvath, Hannum, PhenoAge, and EpiTOC2 (13,14,20,21). The Horvath epigenetic clock is a multi-tissue age predictor, comprised of 353 CpGs, selected via elastic net regression(13). Age acceleration was calculated as the difference between DNA methylation age and chronological age.

DNA methylation levels in repetitive elements were inferred from normalized beta values for each normal gastric mucosa dataset sample using REMP (Repetitive Element Methylation Prediction) version 3.15(22). M-values were extracted using the REMP package, and average M-values were calculated using base R. Statistical analyses were conducted using *limma*(23).

EpiDISH 2.12/HEpiDISH was used to calculate cell type proportions in normal gastric mucosa tissue specimens.(24).We used two reference data sets to infer immune cell type proportions; one reference data from HEpiDISH containing methylation signatures for epithelial, fibroblast, and immune cells, and a second more immune specific reference data set using six immune cell types based on the iterative algorithm Identifying Optimal Libraries (IDOL)(25). Robust partial correlation methods were used.

We identified cell-specific differential methylation using CellDMC(24), a statistical algorithm that uses cell type proportion data, methylation data, and interaction tests in a linear model to identify differential methylation between *H. pylori*-negative controls and *H. pylori* positive infection. CpGs specific to differentially methylated cell types were used to perform eFORGE TF(26) transcription factor analysis using Stomach as reference sample.

## Results

### DNA methylation age and epigenetic aging of gastric mucosa

It has previously been established that DNA methylation (DNAm) undergoes changes as individuals age, and that methylation measures can be used to infer the age of an individual(13). DNA methylation age is an approximation of chronological age and offers the opportunity to calculate age acceleration which is the difference between DNAm age and chronological age. Studies have shown that increased age acceleration is associated with cancer risk(27).

To examine whether *H. pylori* infection is associated with age acceleration, we calculated DNAm age in our study data set **(Table 1)** of 84 samples of normal gastric mucosa using the Horvath epigenetic clock. We used the Horvath epigenetic clock solely to assess epigenetic age and age acceleration due to its multi-tissue applicability making it more accurate for application to gastric mucosae tissue, rather than Hannum and PhenoAge, which were developed using blood samples.

**Table 1.** Cancer and *H. pylori* infection status of subjects from GSE99553

| *H. pylori* Infection Status | Gastric Cancer Status |  |
| --- | --- | --- |
|  | Case (n = 42) | Control (n = 42) |
| Positive (+) | 14 | 14 |
| Past (P) | 14 | 14 |
| Negative (-) | 14 | 14 |

We calculated age acceleration and compared cases with controls, observing significantly higher age acceleration in normal gastric tissue from cancer subjects than controls (P=8.8E-05, **Fig 1A**). When stratifying by cancer status, age acceleration was significantly higher in *H. pylori*-infected normal gastric mucosa of both cancer subjects (P=0.04) and non-cancer controls (P=0.009, **Fig 1B, 1C**). No significant differences in age acceleration were identified between uninfected tissue and tissue with past *H. pylori* infection. These findings indicate that epigenetic aging of gastric tissue is more greatly influenced by cancer status than by *H. pylori* infection status, but that *H. pylori* infection status may contribute to age acceleration of gastric tissue.

**Figure 1.**
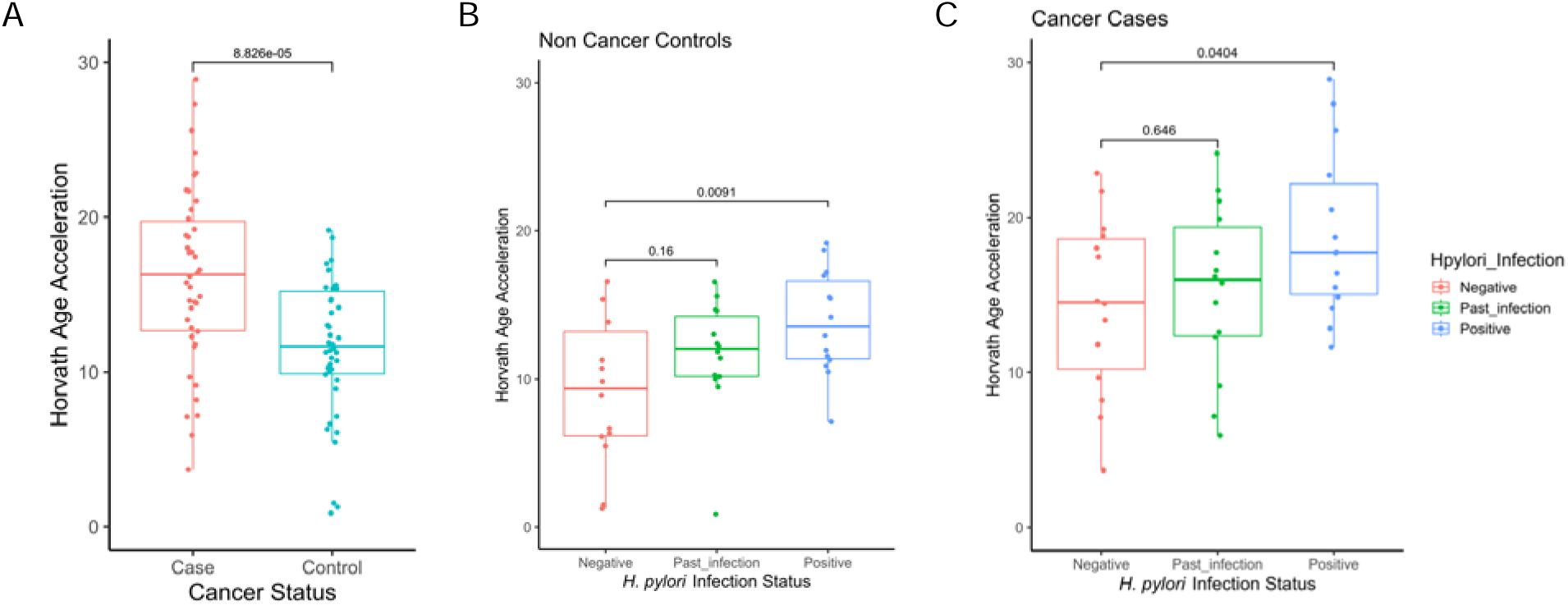
Normal gastric tissue age acceleration by cancer and H. pylori infection status. A) Horvath age acceleration of normal gastric mucosa is higher in subjects with cancer (n=42) than controls (n=42). B) Increased age acceleration in control subject normal gastric mucosa is associated with *H. pylori* infection (n=14), compared with tissue from uninfected controls (n=14). C) In subjects with gastric cancer, normal gastric tissue has increased age acceleration associated with *H. pylori* (n=14) compared with uninfected tissue from cancer cases (n=14).

We next examined whether *H. pylori* infection is associated with the epigenetic mitotic tick rate of normal gastric mucosa using epiTOC2, a mitotic clock that directly estimates the cumulative number of stem cell divisions in tissue(14). The estimated average lifetime intrinsic rate of stem cell division (irS) in the normal gastric mucosa was significantly higher in cancer cases than in controls (P=0.02, **Fig 2A**). Stratifying cancer cases and controls, irS is significantly higher in tissue from subjects with *H. pylori* infection than uninfected tissue in both controls (P=1.5E-06, **Fig 2B**) and in cancer patient normal gastric mucosa (P=1.5E-05, **2C**). Further, past *H. pylori* infection was associated with increased irS in cancer patient normal gastric mucosa compared with uninfected tissue (P=5.0E-05, **Fig 2C**).

**Figure 2.**
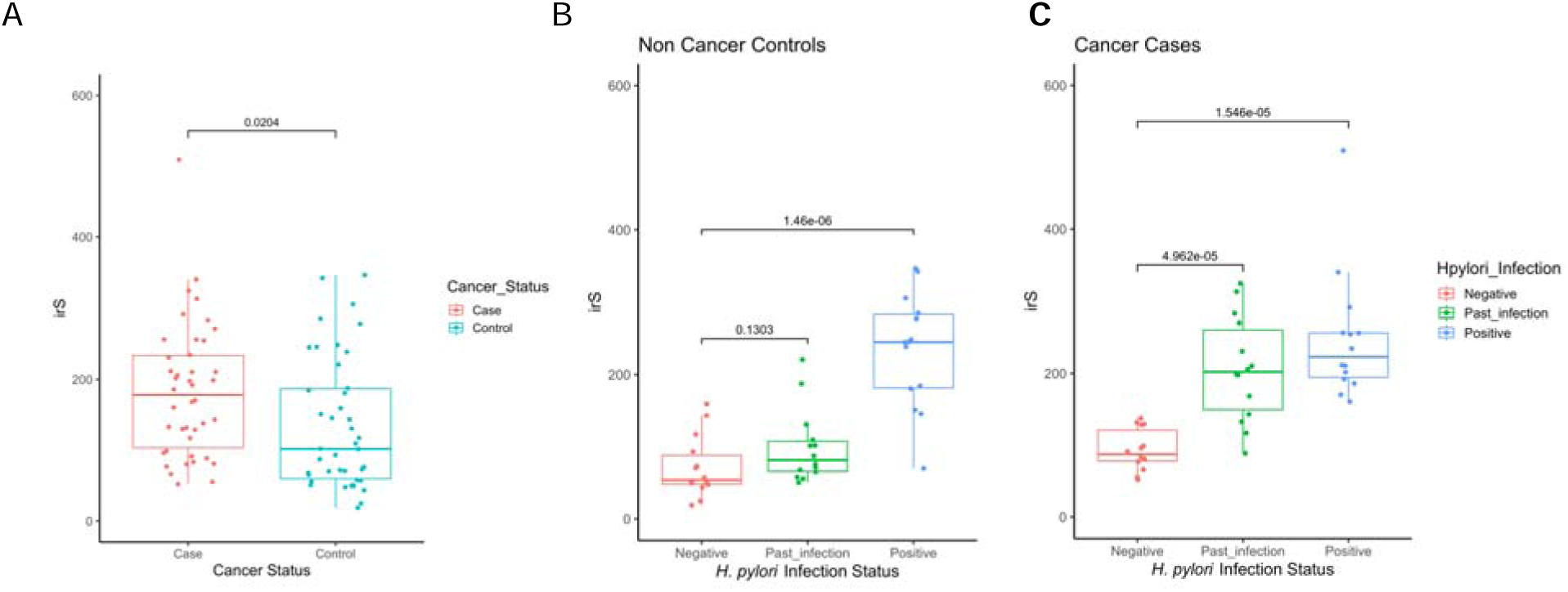
Estimated average lifetime intrinsic rate of stem-cell division (irS) in normal gastric mucosa by cancer and *H. pylori* infection status. **A)** irS was significantly higher in normal gastric mucosa from subjects with gastric cancer (n=42) compared with controls (n-42). **B)** irS was significantly higher in normal gastric mucosae from control subjects with *H. pylori* infection (n=14), compared with tissue from uninfected controls (n=14). **C)** In gastric cancer cases, compared with tissue from uninfected patients (n=14), irS was significantly higher in patients with past *H. pylori* infection (n=14), and in patients with current *H. pylori* infection (n=14).

### *H. pylori* infection status alters gastric tissue epithelial and immune cell type proportions

We assessed cell type proportions using the Hierarchical Epigenetic Dissection of Intra-Sample Heterogeneity (HEpiDISH) algorithm with the IDOL immune reference to infer proportions of eight cell types from DNA methylation data; epithelial, fibroblast, CD8T, CD4T, natural killer (NK), B lymphocytes, monocytes, and neutrophils. The epithelial cell proportions were significantly higher, and fibroblast cell proportions were significantly lower in control subjects’ normal gastric mucosa than in cancer patients’ normal gastric mucosa (**Fig 3A**). In the immune cell component, we observed significantly lower natural killer cells and monocytes in control subjects’ normal gastric mucosa than in cancer patients’ normal gastric mucosa (**Fig 3B**). To assess differences based on *H. pylori* infection status, we investigated cell composition by infection status in control and cancer patient tissues separately. In normal gastric mucosa from controls, we observed significantly lower epithelial cell proportions, higher immune component, and lower fibroblast cell proportions associated with *H. pylori* infection compared with uninfected tissues (**Fig 3C**). Among the cell types comprising the immune component, CD4T, natural killer, B cell, and monocyte cell proportions were all significantly higher in *H. pylori* infected normal gastric mucosa compared with uninfected tissue (**Fig 3D**). Similarly, in tissues from cancer patients, we observed significantly lower epithelial cell and fibroblast cell proportions and higher immune component cell proportions in *H. pylori*-infected normal gastric mucosa compared with uninfected tissue (**Fig 3E**). Among the cell types comprising the immune component in tissues from cancer patients, we observed significantly higher CD4T, natural killer, B cell, and monocyte cell proportions in *H. pylori*-infected normal gastric mucosa compared with uninfected tissue (**Fig 3F**). In tissue from subjects with past *H. pylori* infection, changes in cell proportions were less marked in comparison to *H. pylori* negative cell proportions but demonstrated significantly lower epithelial cell proportions and significantly higher immune cell proportions in tissue from gastric cancer patients with past *H. pylori* infection than in uninfected tissue **(Supplemental Figure 1).**

**Figure 3.**
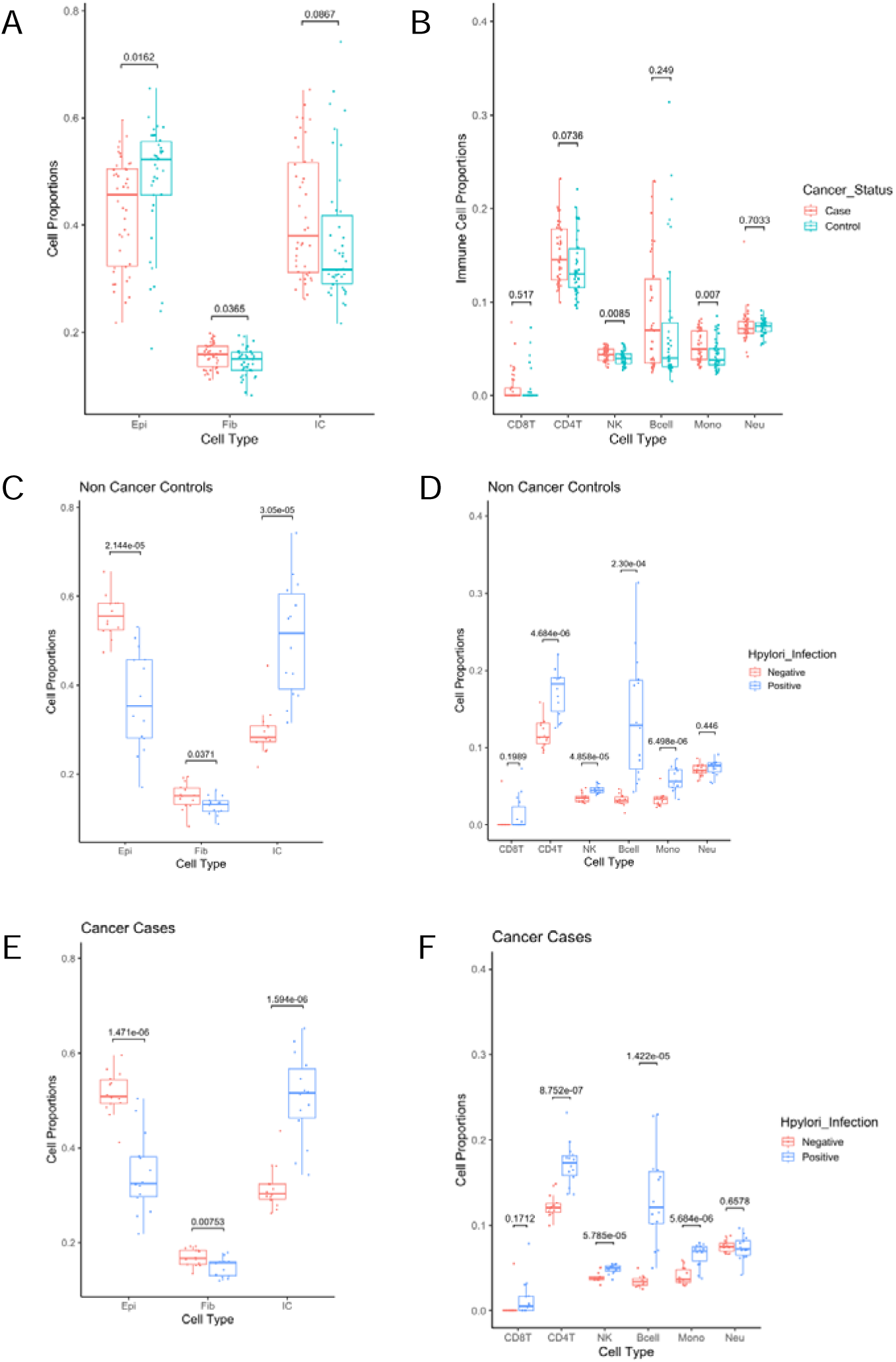
Cell proportions significantly differ by *H. pylori* infection in normal gastric mucosa tissue A) Epithelial cell and Immune Component (IC) proportions demonstrate minor significant differences in epithelial cell and fibroblast cell proportions when comparing normal gastric mucosae from gastric cancer cases to controls. B) Natural killer (NK) and monocyte cell proportions are higher in normal gastric mucosae tissue of cancer cases compared to controls. C) In controls, epithelial cell and fibroblast cell proportions are significantly higher, and IC is significantly lower in HP-normal gastric mucosae compared to HP+. D) The IC in controls is stratified further by specific immune cell component, and significantly higher proportions of CD4T, NK, B cells, and monocytes are observed in normal gastric mucosae from HP+ subjects. E) In gastric cancer cases, epithelial cell and fibroblast cell proportions are significantly higher, and IC is significantly lower in HP-normal gastric mucosae compared to HP+. F) The IC in cases is stratified further by specific immune cell component, and significantly higher CD4T, NK, B cells, and monocytes are observed normal gastric mucosae from HP+ subjects.

**Supplemental Figure 1.**
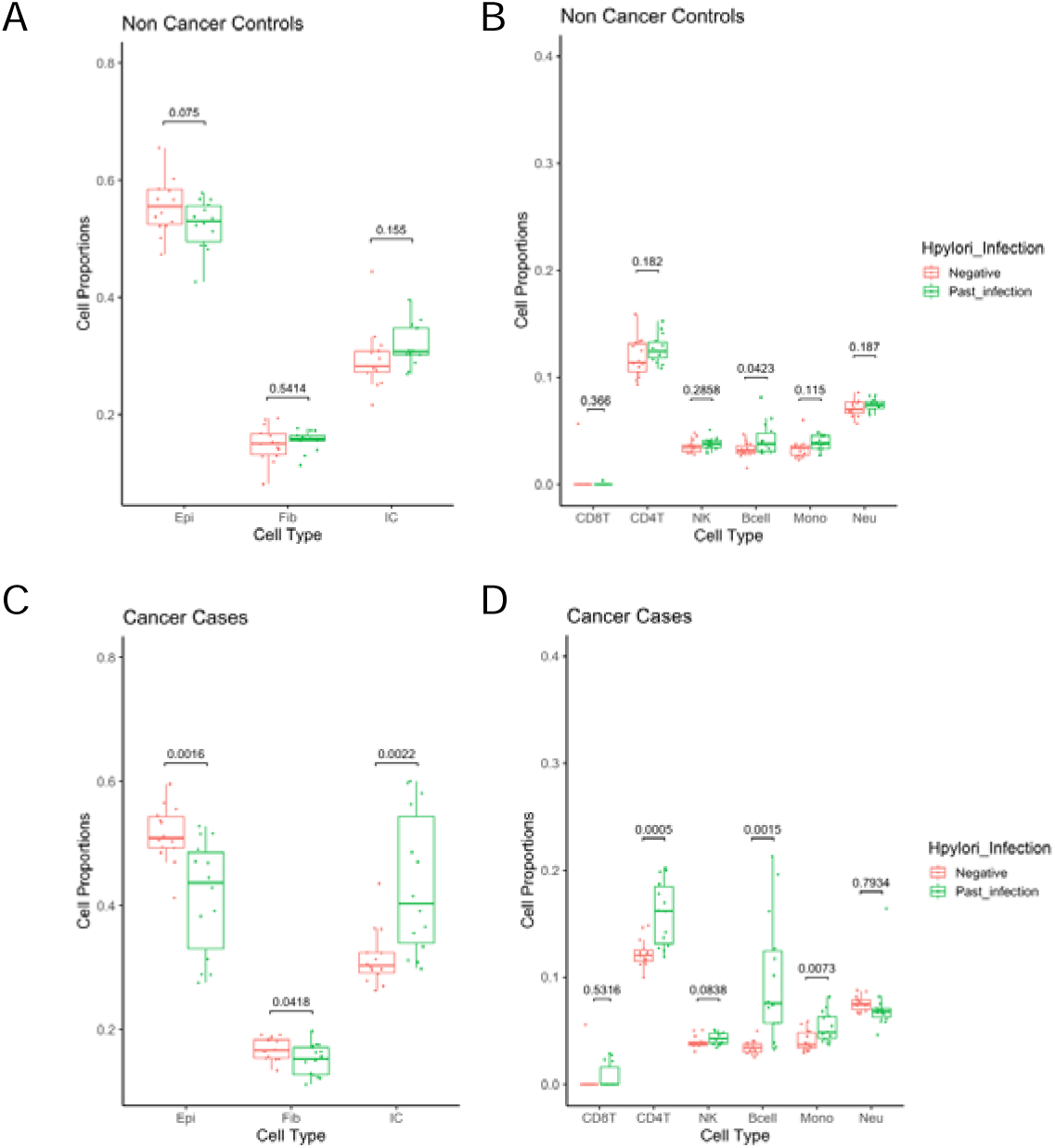
Cell proportions significantly differ in tissue with past *H. pylori* infection in gastric cancer cases. A) Epithelial cell proportions are significantly higher in tissue negative for *H. pylori* compared to past infection in non cancer controls. B) B-cell proportions are significantly higher in tissue with H. pylori past infection compared to H. pylori negative tissue in non cancer controls. C) Epithelial cell proportions are significantly higher and immune component (IC) proportions are significantly lower in tissue negative for *H. pylori* compared to past infection in cancer cases. D) CD4T, B-cell, and Monocyte cell proportions are significantly higher in tissue with *H. pylori* past infection compared to *H. pylori* negative tissue in cancer cases

### Natural killer cells are differentially methylated in normal gastric mucosa from gastric cancer patients with *H. pylori* infection

We next assessed whether certain cell types within the normal gastric mucosa were differentially methylated based on gastric cancer and *H. pylori* infection status. We applied CellDMC(24), using the same HEpiDISH and IDOL cell reference used above. We observed hypermethylation of nine CpG loci in natural killer cells in normal gastric mucosa of gastric cancer patients with positive *H. pylori* infection compared with *H. pylori-*negative tissue (**Table 2**); no other cell types demonstrated significant differential methylation associated with infection. Additionally, no significant differentially methylated cell types were identified in association with *H. pylori* infection in tissues from controls. Using the nine CpG hypermethylated in natural killer cells as input, we performed eFORGE TF (transcription factor) enrichment analysis to investigate the chromatin accessibility signal surrounding the CpG loci and to determine the significance of overlap with transcription factor binding sites. We identified significant overlaps between the input CpG and transcription factor sites, such as at FOXO6, a transcription factor previously shown to be upregulated in gastric cancer tissues and a regulator of cell proliferation in gastric cancer cells(28). Natural killer cells are mediators of protective immunity through critical effector functions of cytotoxicity and production of inflammatory cytokines. Hypermethylation of transcription factor binding sites in natural killer cells may alter the transcription pathways of these cells in the normal gastric mucosa environment in response to *H. pylori* infection, altering their effector functions in a manner which promotes carcinogenesis (**Supplemental Figure 2; Supplemental Table 1**).

**Table 2.**
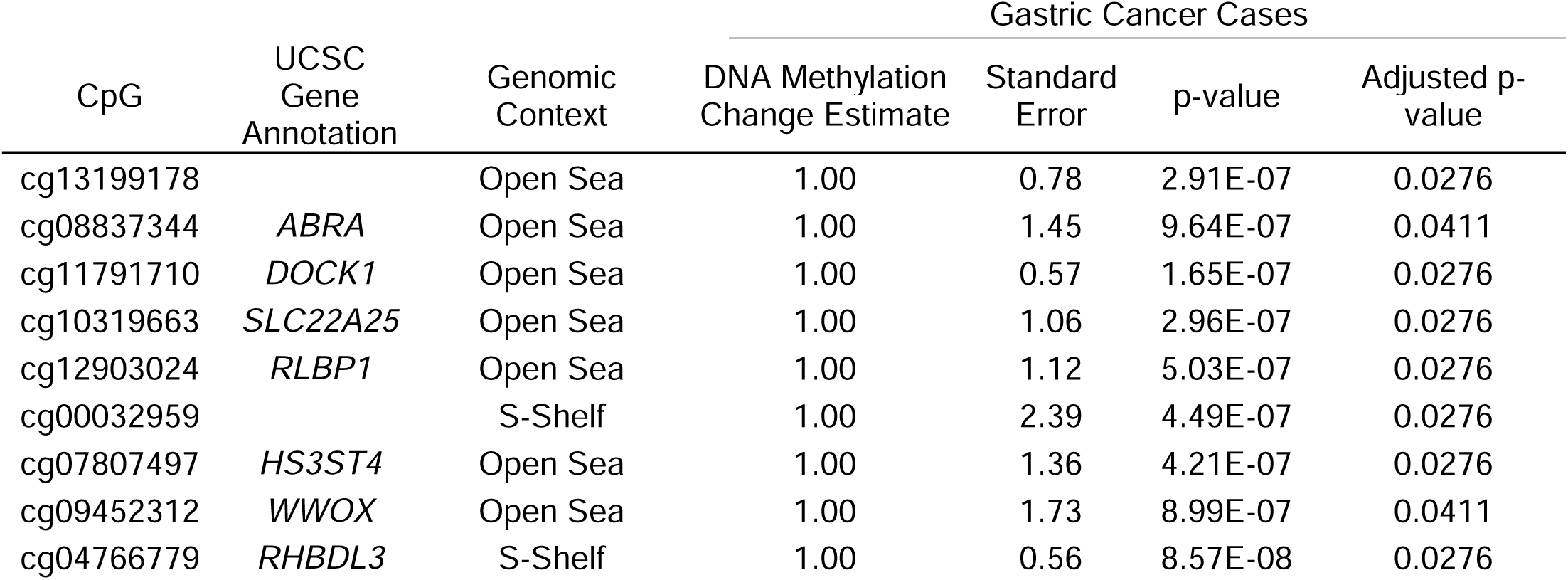
Natural killer cells are differentially methylated in gastric cancer cases with positive *H.pylori* infection but non in controls

**Supplemental Figure 2.**
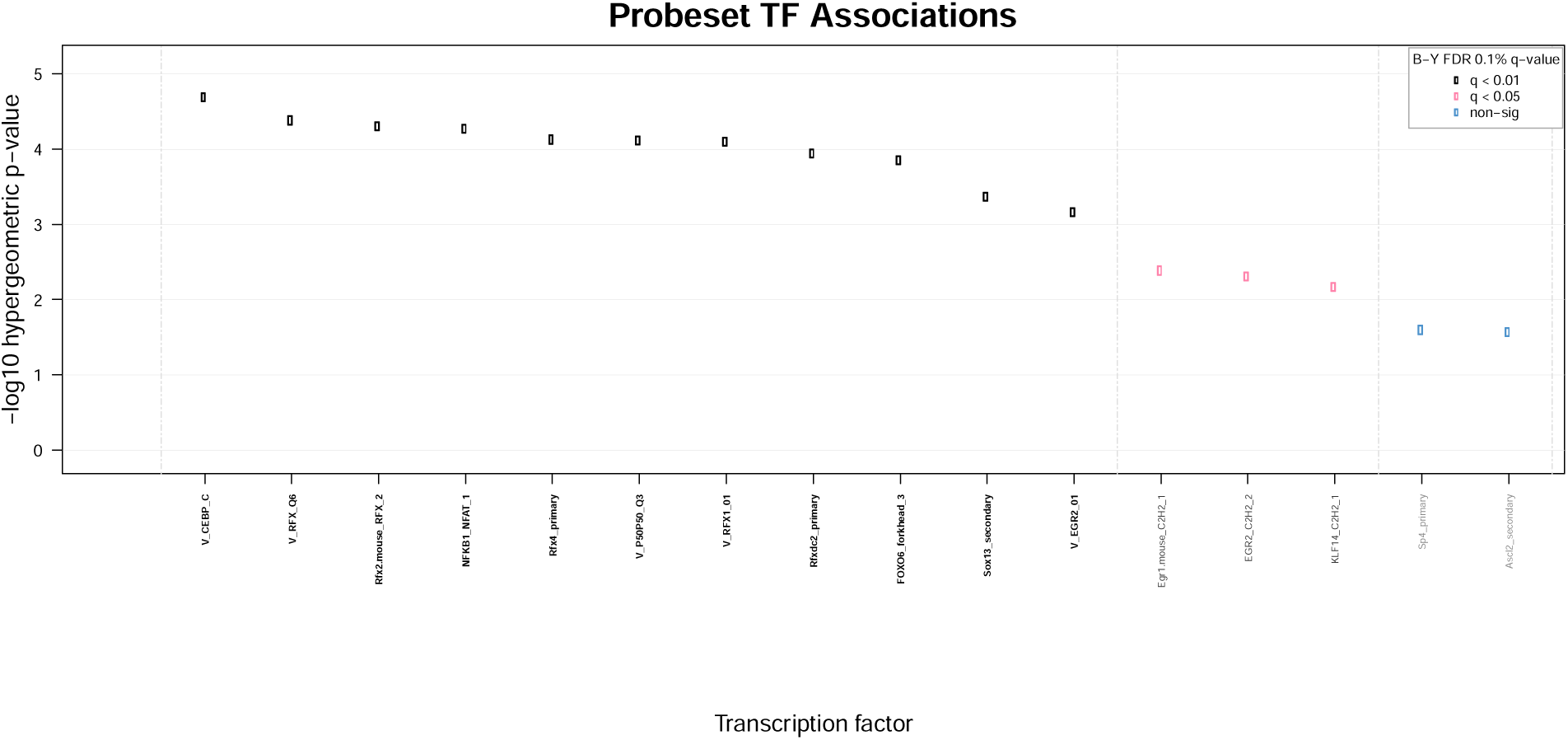
eForge TF Transcription factor analysis, summary plot

**Supplemental Table 1.**
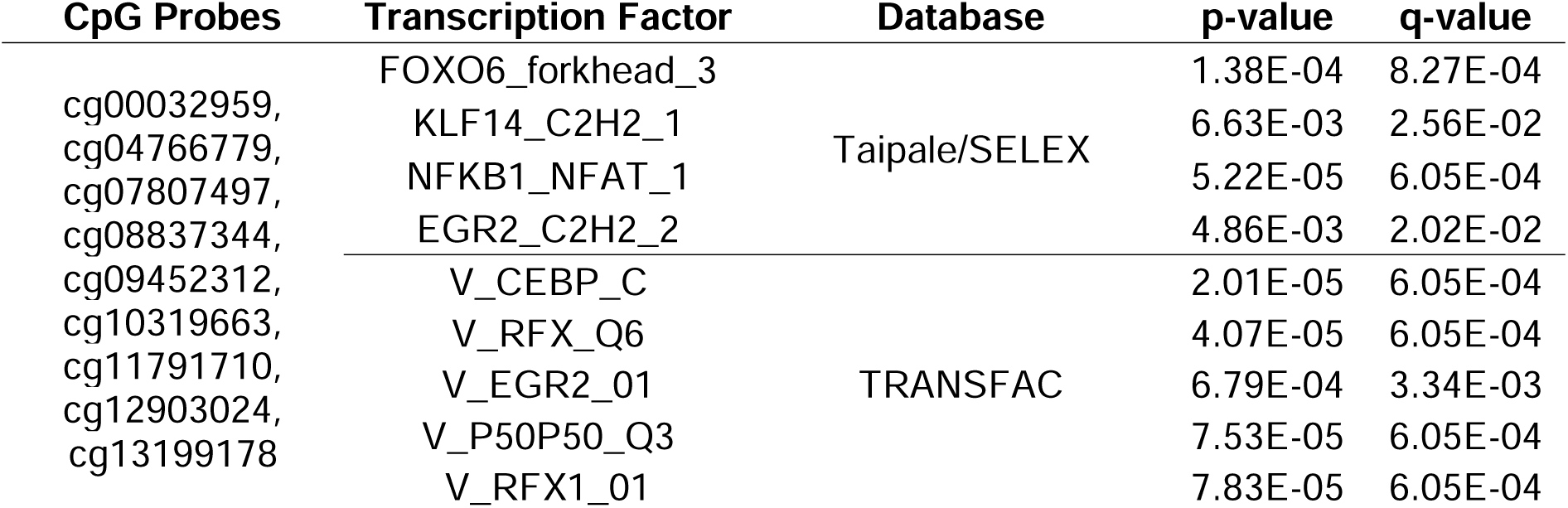
eForge Transcription Factor Analysis Results

### H. pylori infection influences repetitive element methylation

Repetitive elements constitute approximately 50% of the human genome, and short interspersed nuclear elements (SINE) such as *Alu*, long interspersed element-1 (*LINE-1*), and long terminal repeats (*LTR*) within endogenous retroviruses (*ERV*) are epigenetically regulated, and may contribute to variation within cells and tissues of an individual by regulating genes that are located near the targeted repetitive element on the genome(29). Since repetitive elements are mobile and can insert themselves into the genome, they are often hypermethylated as a form of repetitive element regulation. Genomic instability caused by the insertion of repetitive elements may contribute to various diseases, including cancer, through mechanisms such as but not limited to DNA hypomethylation(30). We assessed *H. pylori’s* association with repetitive element methylation in normal gastric mucosa using Repetitive Element Methylation Prediction (REMP) analysis(22), to determine *Alu*, *LINE-1*, and *ERV* methylation. Fitting each repetitive element to a linear model, we investigated the relation of cancer status, *H. pylori* infection status, and cell type as coefficients of interest. Except for a single *Alu* locus, all other repetitive elements were non-significant (**Supplemental Figure 3**). Average methylation of each repetitive element was calculated and *H. pylori* positive infection demonstrated signficantly lower mean *LINE-1* methylation compared to *H. pylori* negative in non cancer controls only (**Supplemental Figure 4).**

**Supplemental Figure 3.**
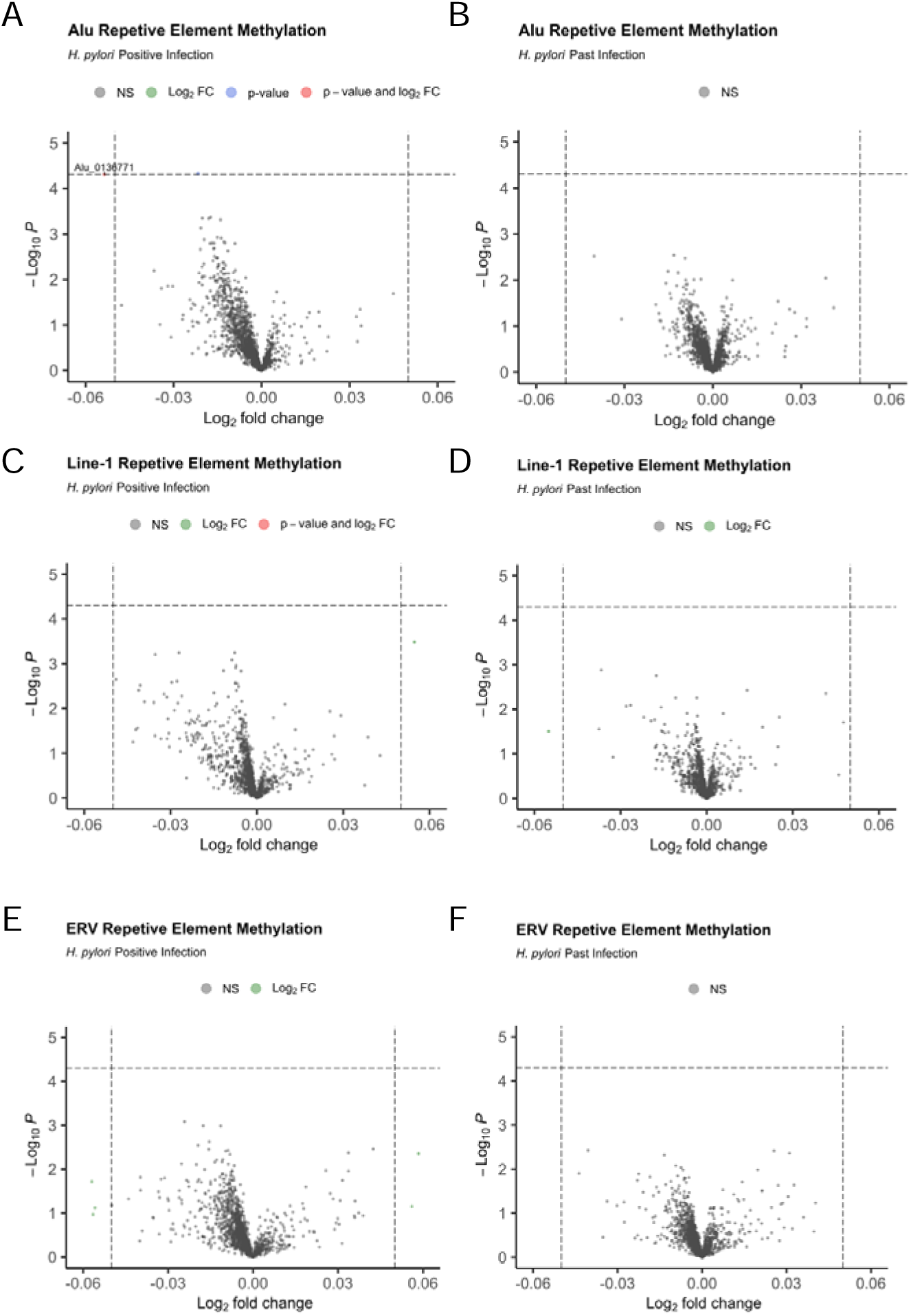
Repetitive Element Methylation Analysis Results. Volcano plots of repetitive element methylation for A) *Alu* methylation and *H. pylori* positive infection, B) *Alu* methylation and *H. pylori* past infection, C) *LINE-1* methylation and *H. pylori* positive infection, D) *LINE-1* methylation and *H. pylori* past infection, E) *ERV* methylation and *H. pylori* positive infection, F) *ERV* methylation and *H. pylori* past infection.

**Supplemental Figure 4.**
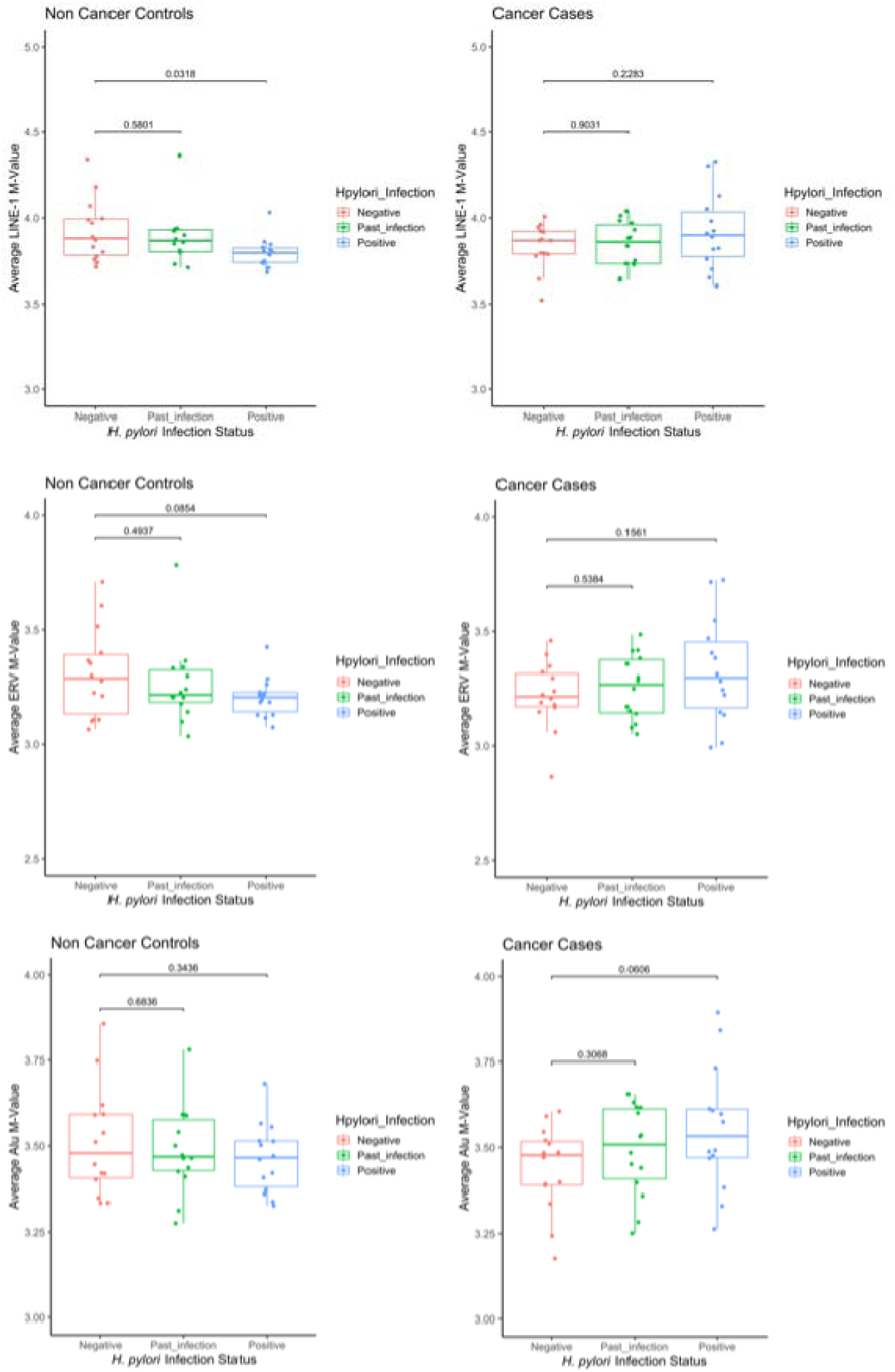
Repetitive Element Average Methylation Average methylation values comparing positive *H. pylori* infection to negative infection, and past *H. pylori* infection to negative infection for A) *LINE-1* methylation in non cancer controls, *LINE-1* methylation in cancer cases, C) *ERV* methylation in non cancer controls, D) *ERV* methylation in cancer cases, E) *Alu* methylation in non cancer controls, and F) *Alu* methylation in cancer cases.

## Discussion

This study presents an analysis of DNA methylation in normal gastric mucosa of gastric cancer cases and control subjects with *H. pylori* infection, uninfected, and past infection, and details changes in epigenetic changes based on *H. pylori*-positive infection status, independent of cancer status.

Using epigenetic clocks to statistically assess the epigenetic age and age acceleration of the normal gastric mucosae, we identified differences in the intrinsic rate of stem cell division per sample, specifically that intrinsic rates of stem cell division are highest in subjects with a positive *H. pylori* infection in both cancer cases and controls, indicating increased mitotic age of those normal gastric mucosae. Tissue mitotic age has been linked to the risk of neoplastic transformation in tissues and is directly influenced by cell turnover rate(15). Our findings indicate that there is a greater rate of cell division in HP+ normal gastric mucosae of both cancer cases and non-cancer controls compared to HP-, suggesting that *H. pylori* is influencing the induction of cell proliferation and that cell proliferation is not being driven by cancer case status alone. It has been previously established that induction of cell proliferation is driven by bacterial infection and chronic inflammation, such as with *H. pylori*(31). Considering induction of cell proliferation is observed in normal gastric mucosae of HP+ subjects in both cancer cases and non-cancer controls, we can posit that infection via *H. pylori* induces changes in normal tissues that may contribute to an environment that is more susceptible to carcinogenesis due to increased cell proliferation and chronic inflammation.

To better understand how normal gastric mucosae of both gastric cancer cases and controls may be altered by *H. pylori* infection, we assessed epithelial, fibroblast, and immune cell proportions and observed significantly lower epithelial cell, lower fibroblast, and higher immune cell component proportions in HP+ compared to HP-normal gastric mucosae in both gastric cancer cases and controls. More specifically, we identified significantly higher CD4T, natural killer, B-cell, and monocyte cell proportions in HP+ tissue compared to HP-. The observed differences in B-cell populations are of particular interest, considering the role of B-cells in gastric mucosa-associated lymphoid tissue (MALT) lymphoma and considering previous studies have demonstrated incidence of gastric MALT lymphoma is dependent on *H. pylori* infection rate(32). Infection with H. pylori of normal gastric mucosa leads to the development of organized lymphoid tissue within the gastric mucosa, and the accumulation of lymphoid tissue containing B-cell follicles(33). In our analysis of cell type proportion differences by *H. pylori* infection status, B-cell proportions were significantly higher in HP+ normal gastric mucosa than in HP-, indicating changes to B-cell proportions occurring within the normal gastric mucosa based on infection status, independent of cancer status. Eradication of *H. pylori* as indicated by a past infection demonstrates significant differences in B-cell proportions, but not as significant as HP+ infection, highlighting how *H. pylori* eradication may allow the normal gastric mucosae to return to a non-carcinogenic environment, while prolonged positive H. pylori infection may contribute to changes within the normal gastric mucosa that are conducive to carcinogenesis.

Additionally, we observed the starkest difference in cell proportions of CD4T and natural killer cells between HP+ and HP-samples in both gastric cancer cases and controls. Previous studies have shown higher levels of CD4T cells in *H. pylori-*infected gastric mucosae produce high quantities of interferon-gamma (IFN-γ) and low levels of interleukin-4 (IL4), which together may be contributors to gastritis due to increased inflammation from increased cytokine levels(34). Natural killer cell activity has previously been correlated to clinical stage, lymphatic and vascular invasion, lymph node metastases, and prognosis in gastric cancer patients, with natural killer cell number and activity found to be decreased with the progression of gastric cancer.(35) However, in non-cancer gastric mucosa infected with *H. pylori*, there is evidence that natural killer cells are activated directly by *H. pylori* antigens and by *H. pylori* bacteria and that this activation induces the production of IFN-γ by natural killer cells(36). Higher proportions of natural killer cells in HP+ gastric mucosa are consistent biologically considering the role of natural killer cells in the innate immune response; however, hypermethylation of the genome in these cell types, as indicated in our analyses, may indicate specific cell programs may be transcriptionally repressed in natural killer cells. Our enrichment analysis on the hypermethylated CpGs identified in natural killer cells demonstrated significant enrichment of transcription binding motifs including FOXO6 and NF-κB1/NFAT. FOX transcription factors bind chromatin and can act as co-activators and transcriptional repressors, and contribute to cancer at various levels of regulation(37). NF-κB1 and NFAT dimers regulate the transcription of target genes for T-lymphocyte activation, playing a crucial role in the regulation of cell growth and apoptosis(38). Hypermethylation of these transcription factor motifs may affect binding of specific transcription factors, altering the transcriptional pathways of natural killer cells. Recently, a study demonstrated infection with *H. pylori* resulted in hypermethylation of the promotor region of the *SOCS-1* gene in both tumor tissue and normal adjacent tissue from gastric cancer patients infected with *H. pylori* (39), a gene that suppresses cytokine signaling after induction by high levels of cytokines including IFN-γ. A potential pathway to carcinogenesis exacerbated by *H. pylori* in gastric mucosae may be a combined effect of increased CD4T cells in HP+ tissue and increased natural killer cells in HP+ tissue, which both increase levels of IFN-γ. IFN-γ has been identified as a critical promoter of cell atrophy and progression from gastritis to gastric atrophy and metaplasia in mouse models(40), and high production levels of IFN-γ can lead to prolonged inflammation(41).

Limitations in our research include a small sample size, which may not be powerful enough to detect signals when assessing repetitive elements or sufficient to uncover subtler differences when assessing tissue with a past *H. pylori* infection. Additionally, while data used within this analysis was collected from normal gastric mucosa as indicated in the original study, the tissue may still be tumor-adjacent and/or tumor influenced, further influencing or impacting any weaker signals in our analyses. Infection with *H. pylori* is marked by damage to the gastric epithelia via bacterial colonization, and it is reasonable that epithelial and fibroblast proportions are lowest in *H. pylori*-positive subjects. Additionally, CD4T, NK, B lymphocyte, and monocyte cell proportions are highest in subjects with a positive *H. pylori* infection in both cases and controls, indicating immune cell recruitment and activity within *H. pylori*-positive gastric tissues. *H. pylori*-induced inflammation involves activation of the host immune system and infiltration of immune cells(42), and increased B cell proportions may rationalize increased CD4T cell proportions in *H. pylori*-positive subjects since B cells are necessary for optimal T cell activation and immunity(43).

Our work provides a basis for further assessing cell heterogeneity within gastric cancer and assessing how *H. pylori* may influence specific cell populations, potentially influencing certain cell types as drivers of carcinogenesis. Our findings provide further evidence of gastric cancer molecular mechanisms, marked by changes to DNA methylation, further exacerbated by *H. pylori* infection.

## Declarations

### Ethics approval and consent to participate

Not Applicable

### Consent for publication

Not Applicable

### Availability of data and materials

Data used in this study are publically available in the GEO repository (GSE99553).

### Funding

R01CA253976

R01CA216265

P20GM104416

P30CA023108

CDMRP/Department of Defense (W81XWH-20-1-0778)

### Authors’ contributions

IMV and LAS designed the study. IMV performed the analyses and wrote the manuscript. All authors revised the manuscript. All authors read and approved the final manuscript.

## Acknowledgements

Not Applicable

## References

1. Tan P, Yeoh KG. Genetics and Molecular Pathogenesis of Gastric Adenocarcinoma. Gastroenterology. 2015 Oct 1;149(5):1153–1162.e3.

2. Alipour M. Molecular Mechanism of Helicobacter pylori-Induced Gastric Cancer. J Gastrointest Canc. 2021 Mar 1;52(1):23–30.

3. Sung H, Ferlay J, Siegel RL, Laversanne M, Soerjomataram I, Jemal A, et al. Global Cancer Statistics 2020: GLOBOCAN Estimates of Incidence and Mortality Worldwide for 36 Cancers in 185 Countries. CA: A Cancer Journal for Clinicians. 2021;71(3):209–49.

4. Kumar S, Metz DC, Ellenberg S, Kaplan DE, Goldberg DS. Risk Factors and Incidence of Gastric Cancer After Detection of Helicobacter pylori Infection: A Large Cohort Study. Gastroenterology. 2020 Feb 1;158(3):527–536.e7.

5. Chung HW, Lim JB. Role of the tumor microenvironment in the pathogenesis of gastric carcinoma. World J Gastroenterol. 2014 Feb 21;20(7):1667–80.

6. Yamashita S, Nanjo S, Rehnberg E, Iida N, Takeshima H, Ando T, et al. Distinct DNA methylation targets by aging and chronic inflammation: a pilot study using gastric mucosa infected with Helicobacter pylori. Clin Epigenet. 2019 Dec;11(1):191.

7. Berdasco M, Esteller M. Aberrant Epigenetic Landscape in Cancer: How Cellular Identity Goes Awry. Developmental Cell. 2010 Nov 16;19(5):698–711.

8. Ehrlich M. DNA methylation in cancer: too much, but also too little. Oncogene. 2002 Aug;21(35):5400–13.

9. Nishiyama A, Nakanishi M. Navigating the DNA methylation landscape of cancer. Trends in Genetics. 2021 Nov 1;37(11):1012–27.

10. Robertson KD. DNA methylation and human disease. Nat Rev Genet. 2005 Aug;6(8):597–610.

11. Niwa T, Tsukamoto T, Toyoda T, Mori A, Tanaka H, Maekita T, et al. Inflammatory Processes Triggered by *Helicobacter pylori* Infection Cause Aberrant DNA Methylation in Gastric Epithelial Cells. Cancer Research. 2010 Feb 15;70(4):1430– 40.

12. Pappalardo XG, Barra V. Losing DNA methylation at repetitive elements and breaking bad. Epigenetics & Chromatin. 2021 Jun 3;14(1):25.

13. Horvath S. DNA methylation age of human tissues and cell types. Genome Biology. 2013 Dec 10;14(10):3156.

14. Teschendorff AE. A comparison of epigenetic mitotic-like clocks for cancer risk prediction. Genome Medicine. 2020 Jun 24;12(1):56.

15. Tomasetti C, Vogelstein B. Variation in cancer risk among tissues can be explained by the number of stem cell divisions. Science. 2015 Jan 2;347(6217):78–81.

16. Woo HD, Fernandez-Jimenez N, Ghantous A, Degli Esposti D, Cuenin C, Cahais V, et al. Genome-wide profiling of normal gastric mucosa identifies Helicobacter pylori- and cancer-associated DNA methylome changes. Int J Cancer. 2018 Aug 1;143(3):597–609.

17. Xu Z, Niu L, Li L, Taylor JA. ENmix: a novel background correction method for Illumina HumanMethylation450 BeadChip. Nucleic Acids Res. 2016 Feb 18;44(3):e20.

18. Teschendorff AE, Marabita F, Lechner M, Bartlett T, Tegner J, Gomez-Cabrero D, et al. A beta-mixture quantile normalization method for correcting probe design bias in Illumina Infinium 450 k DNA methylation data. Bioinformatics. 2013 Jan 15;29(2):189–96.

19. Zhou W, Laird PW, Shen H. Comprehensive characterization, annotation and innovative use of Infinium DNA methylation BeadChip probes. Nucleic Acids Res. 2017 Feb 28;45(4):e22.

20. Hannum G, Guinney J, Zhao L, Zhang L, Hughes G, Sadda S, et al. Genome-wide methylation profiles reveal quantitative views of human aging rates. Mol Cell. 2013 Jan 24;49(2):359–67.

21. Levine ME, Lu AT, Quach A, Chen BH, Assimes TL, Bandinelli S, et al. An epigenetic biomarker of aging for lifespan and healthspan. Aging (Albany NY). 2018 Apr 17;10(4):573–91.

22. Zheng Y, Joyce BT, Liu L, Zhang Z, Kibbe WA, Zhang W, et al. Prediction of genome-wide DNA methylation in repetitive elements. Nucleic Acids Res. 2017 Sep 6;45(15):8697–711.

23. Ritchie ME, Phipson B, Wu D, Hu Y, Law CW, Shi W, et al. limma powers differential expression analyses for RNA-sequencing and microarray studies. Nucleic Acids Research. 2015 Apr 20;43(7):e47–e47.

24. Zheng SC, Breeze CE, Beck S, Teschendorff AE. Identification of differentially methylated cell types in epigenome-wide association studies. Nat Methods. 2018 Dec;15(12):1059–66.

25. Salas LA, Koestler DC, Butler RA, Hansen HM, Wiencke JK, Kelsey KT, et al. An optimized library for reference-based deconvolution of whole-blood biospecimens assayed using the Illumina HumanMethylationEPIC BeadArray. Genome Biol. 2018 May 29;19:64.

26. Breeze CE, Reynolds AP, van Dongen J, Dunham I, Lazar J, Neph S, et al. eFORGE v2.0: updated analysis of cell type-specific signal in epigenomic data. Bioinformatics. 2019 Nov 15;35(22):4767–9.

27. Hong C, Yang S, Wang Q, Zhang S, Wu W, Chen J, et al. Epigenetic Age Acceleration of Stomach Adenocarcinoma Associated With Tumor Stemness Features, Immunoactivation, and Favorable Prognosis. Frontiers in Genetics [Internet]. 2021 [cited 2022 Jun 7];12. Available from: https://www.frontiersin.org/article/10.3389/fgene.2021.563051

28. Qinyu L, Long C, Zhen-dong D, Min-min S, Wei-ze W, Wei-ping Y, et al. FOXO6 promotes gastric cancer cell tumorigenicity via upregulation of C-myc. FEBS Letters. 2013 Jul 11;587(14):2105–11.

29. Slotkin RK, Martienssen R. Transposable elements and the epigenetic regulation of the genome. Nat Rev Genet. 2007 Apr;8(4):272–85.

30. Kong Y, Rose CM, Cass AA, Williams AG, Darwish M, Lianoglou S, et al. Transposable element expression in tumors is associated with immune infiltration and increased antigenicity. Nat Commun. 2019 Nov 19;10(1):5228.

31. Ushijima T, Hattori N. Molecular Pathways: Involvement of Helicobacter pylori– Triggered Inflammation in the Formation of an Epigenetic Field Defect, and Its Usefulness as Cancer Risk and Exposure Markers. Clinical Cancer Research. 2012 Feb 14;18(4):923–9.

32. Doglioni C, Moschini A, de Boni M, Wotherspoon AC, Isaacson PG. High incidence of primary gastric lymphoma in northeastern Italy. The Lancet. 1992 Apr 4;339(8797):834–5.

33. Du MQ, Isaccson PG. Gastric MALT lymphoma: from aetiology to treatment. The Lancet Oncology. 2002 Feb;3(2):97–104.

34. Chiba T, Marusawa H, Seno H, Watanabe N. Mechanism for gastric cancer development by Helicobacter pylori infection. Journal of Gastroenterology and Hepatology. 2008;23(8pt1):1175–81.

35. Du Y, Wei Y. Therapeutic Potential of Natural Killer Cells in Gastric Cancer. Front Immunol. 2019 Jan 21;9:3095.

36. Yun CH, Lundgren A, Azem J, Sjöling Å, Holmgren J, Svennerholm AM, et al. Natural Killer Cells and Helicobacter pylori Infection: Bacterial Antigens and Interleukin-12 Act Synergistically To Induce Gamma Interferon Production. Infection and Immunity. 2005 Mar;73(3):1482–90.

37. Bach DH, Long NP, Luu TTT, Anh NH, Kwon SW, Lee SK. The Dominant Role of Forkhead Box Proteins in Cancer. International Journal of Molecular Sciences. 2018 Oct;19(10):3279.

38. Arlt A, Schäfer H, Kalthoff H. The ‘N-factors’ in pancreatic cancer: functional relevance of NF-κB, NFAT and Nrf2 in pancreatic cancer. Oncogenesis. 2012 Nov;1(11):e35–e35.

39. Jan I, Rather RA, Mushtaq I, Malik AA, Besina S, Baba AB, et al. Helicobacter pylori Subdues Cytokine Signaling to Alter Mucosal Inflammation via Hypermethylation of Suppressor of Cytokine Signaling 1 Gene During Gastric Carcinogenesis. Frontiers in Oncology [Internet]. 2021 [cited 2023 Apr 20];10. Available from: https://www.frontiersin.org/articles/10.3389/fonc.2020.604747

40. Osaki LH, Bockerstett KA, Wong CF, Ford EL, Madison BB, DiPaolo RJ, et al. Interferon-γ directly induces gastric epithelial cell death and is required for progression to metaplasia. J Pathol. 2019 Apr;247(4):513–23.

41. Schoenborn JR, Wilson CB. Regulation of Interferon-γ During Innate and Adaptive Immune Responses. In: Advances in Immunology [Internet]. Academic Press; 2007 [cited 2023 Apr 20]. p. 41–101. Available from: https://www.sciencedirect.com/science/article/pii/S0065277607960022

42. Semper RP, Vieth M, Gerhard M, Mejías-Luque R. Helicobacter pylori Exploits the NLRC4 Inflammasome to Dampen Host Defenses. The Journal of Immunology. 2019 Oct 15;203(8):2183–93.

43. DiLillo DJ, Yanaba K, Tedder TF. B cells are required for optimal CD4+ and CD8+ T cell tumor immunity: therapeutic B cell depletion enhances B16 melanoma growth in mice. J Immunol. 2010 Apr 1;184(7):4006–16.

